# Systematic functional dissection of germline noncoding risk variants impacting clonal hematopoiesis

**DOI:** 10.64898/2026.01.21.700897

**Authors:** Trieu Nguyen, Jessica Jeejan, Takeshi Iwasaki, Susan Kales, Joyeeta Chakraborty, Chie Yanase, Aruna Shekhar, Dominika Kwasniak, Aditi Hegde, Richard Voit, Kristy Stengel, Keisuke Ito, Ryan Tewhey, Satish Nandakumar

## Abstract

Clonal hematopoiesis of indeterminate potential (CHIP) is a precursor condition characterized by the expansion of blood cell clones harboring somatic mutations originating in hematopoietic stem cells (HSCs). Since individuals with CHIP face a high risk of developing myeloid malignancies, targeting CHIP clones could provide a viable strategy for leukemia prevention. Despite its clinical significance, the mechanisms underlying CHIP predisposition and progression remain poorly understood. Recent genome wide association studies (GWAS) have identified several non-coding genetic loci that are strongly associated with CHIP; however, their underlying mechanisms still remain unknown. We hypothesize that risk variants in these non-coding loci modulate enhancer elements active in HSCs. To test this, we selected 1,374 non-coding variants from 51 loci associated for CHIP risk in the UK Biobank and screened them for regulatory activity using a Massively Parallel Reporter Assay (MPRA). We performed our lentiviral MPRA screen in MUTZ-3 cells, a human hematopoietic cell line relevant to HSCs, which express CD34 surface marker and are dependent on HSC-specific transcription factors. Using a MPRA library of ∼73,000 constructs in CD34+ fraction of MUTZ-3 cells, we identified 87 variants representing 32 GWAS loci. We used targeted genome editing to demonstrate endogenous enhancer activity across 3 MPRA variants that affect the transcription of NKD2, FLT3, and MSI2. Our functional studies on MSI2 indicate that presence of higher levels of MSI2 mediated by CHIP risk allele enhances the clonal expansion of TET2 knockout hematopoietic stem and progenitor cells, providing a mechanistic link whereby non-coding genetic variants can influence the expansion of mutant CHIP clones.

## Introduction

Aging is accompanied with gradual accumulation of somatic mutations across various tissues in the human body^1^. In the hematopoietic system, mutations that confer fitness advantage to the hematopoietic stem cell (HSC) can lead to clonal expansion of mutant HSCs and their progeny in otherwise healthy individuals, a phenomenon termed clonal hematopoiesis of indeterminate potential (CHIP)^2^. CHIP becomes more prevalent with age and is estimated to affect ∼10% of the population older than 70 years of age^3,4^. CHIP is associated with an increased risk of hematologic malignancies and a range of nonhematologic conditions, including cardiovascular disease and chronic inflammatory states. Despite its importance, we still don’t understand the mechanisms of CHIP predisposition and its progression. Lineage tracing studies estimate that CHIP clones arise decades earlier before detection and sometimes as early as during birth indicating that CHIP driver mutations are insufficient to fully drive clonal expansion and may require additional factors^5,6^.

Recent GWAS studies have shown that germline genetic background influences the propensity to develop CHIP. The NHLBI TOPMed study containing 4,229 CHIP carriers from different ancestries identified three genetic risk loci^7^. Two recent GWAS studies based on UK Biobank with further expanded cohort sizes containing ∼10,000 and ∼40,000 CHIP carriers have identified 14 and 24 genetic risk loci respectively^8,9^. However, specific causal variants remain unknown due to linkage disequilibrium and the prevalence of variants in non-coding regions, which make it difficult to distinguish causal alleles from bystanders. In previous work, we and others have directly demonstrated that at particular loci, modulation of gene regulatory elements in HSCs underlies genetic risk for myeloid malignancies. For example, we identified a germline CHIP variant that inactivates a TET2 enhancer in human hematopoietic stem and progenitor cells (HSPCs), and another variant at a GFI1B enhancer linked to JAK2^V617F^ CHIP and myeloproliferative neoplasms that influences HSC expansion^7,10^. Similarly, another CHIP risk variant in the TCL1A promoter was shown to influence gene expression on CHIP mutant HSCs^11^.

Here, we now utilized a high-throughput assay termed massively parallel reporter assay (MPRA) to systematically measure enhancer activity of 1,374 non-coding variants within 51 fine-mapped loci from a comprehensive GWAS study of CHIP risk. Previously MPRA approach was only feasible in cell lines due to limitations in delivery of the reporter library^12,13^. Now we utilized a lentiviral based integrating version of the MPRA (LentiMPRA) to evaluate CHIP risk variants directly for the first time in HSPC relevant cell populations^14^. Our MPRA identified 87 functional variants representing 32 (63%) of the original 51 fine-mapped loci (median of 1 variant/loci). We used CRISPR/Cas9 genome editing to confirm the endogenous activity of three variant harboring regulatory elements (at MSI2, TERT, FLT3 loci). Follow-up studies of the target gene MSI2, demonstrated its critical role in promoting the clonal expansion of TET2-mutant hematopoietic stem and progenitor cells (HSPCs), thereby establishing a mechanistic link through which non-coding genetic variants can drive the selective advantage and outgrowth of mutant CHIP clones.

## Results

To select the non-coding CHIP risk variants for LentiMPRA, we focused on the largest genome wide association study (GWAS) to date on clonal hematopoiesis. This study was conducted on 628,388 individuals and identified 24 genetic loci associated with CHIP. Bayesian fine mapping analysis using FINAMAP tool identified 51 conditionally independent signals containing 3842 variants associated with CHIP^9^. We overlapped all fine-mapped variants to ATAC-seq peaks across 18 human hematopoietic stem and progenitor cell populations, resulting in identifying 560 CHIP risk variants that we prioritized for MPRA **(Fig. S1A)**. We also included 127 variants that has high posterior probability (PP> 0.05) but did not overlap with ATAC-seq peaks (non-ATAC variants). Both the risk and the non-risk allele were tested for all 687 variants resulting in 1,374 constructs **(Fig. S1A)**. In addition, we also included 200 negative controls that are in the coding regions of the human genome (ORF controls) and showed minimal or low regulatory activity in previous studies^15^. The library also had 39 positive controls which included validated HSPC enhancers from previous studies. We included two enhancers that control GATA2, a critical regulator of HSCs, the GATA2 oncogenic enhancer at 3q21 that is commonly rearranged in inv(3) AML and the GATA2 +9.5 intronic enhancer that is mutated in GATA2 haploinsufficiency syndromes^16–18^. Additionally, we tested the PU.1 upstream regulatory element (URE) that regulates PU.1 expression in HSCs^19^ and a CHIP risk variant containing TET2 enhancer that we validated in our previous work^7^. This resulted in a final library size of 1613 constructs for our MPRA experiment **(Fig. S1A, Table S1)**.

We modified a recently developed lentiviral MPRA to screen for medium sized oligonucleotide fragments (250bp) in an integrated format^14^. For each allele (major/minor), we synthesized constructs containing CHIP risk variant in the center, with surrounding genomic region, 125bp upstream and downstream of the variant. The oligos were cloned upstream of the minimal promoter and 20 bp barcode attached **(Fig 1A)**. After cloning, we sequenced the plasmid library to saturation to determine the enhancer-barcode association using NextSeq Illumina sequencing. From the 13 million sequencing reads that were obtained, 749,801 distinct barcode-enhancer associations were identified **(Fig. S1B)**. A total of 1588/ 1613 enhancers were captured with a 2% failure rate which is consistent with our previous experience in generating similar libraries. Sequences were uniformly distributed within the library and paired with a median of around 305 of barcodes per oligo **(Fig. S1C,D)**.

**Figure 1.**
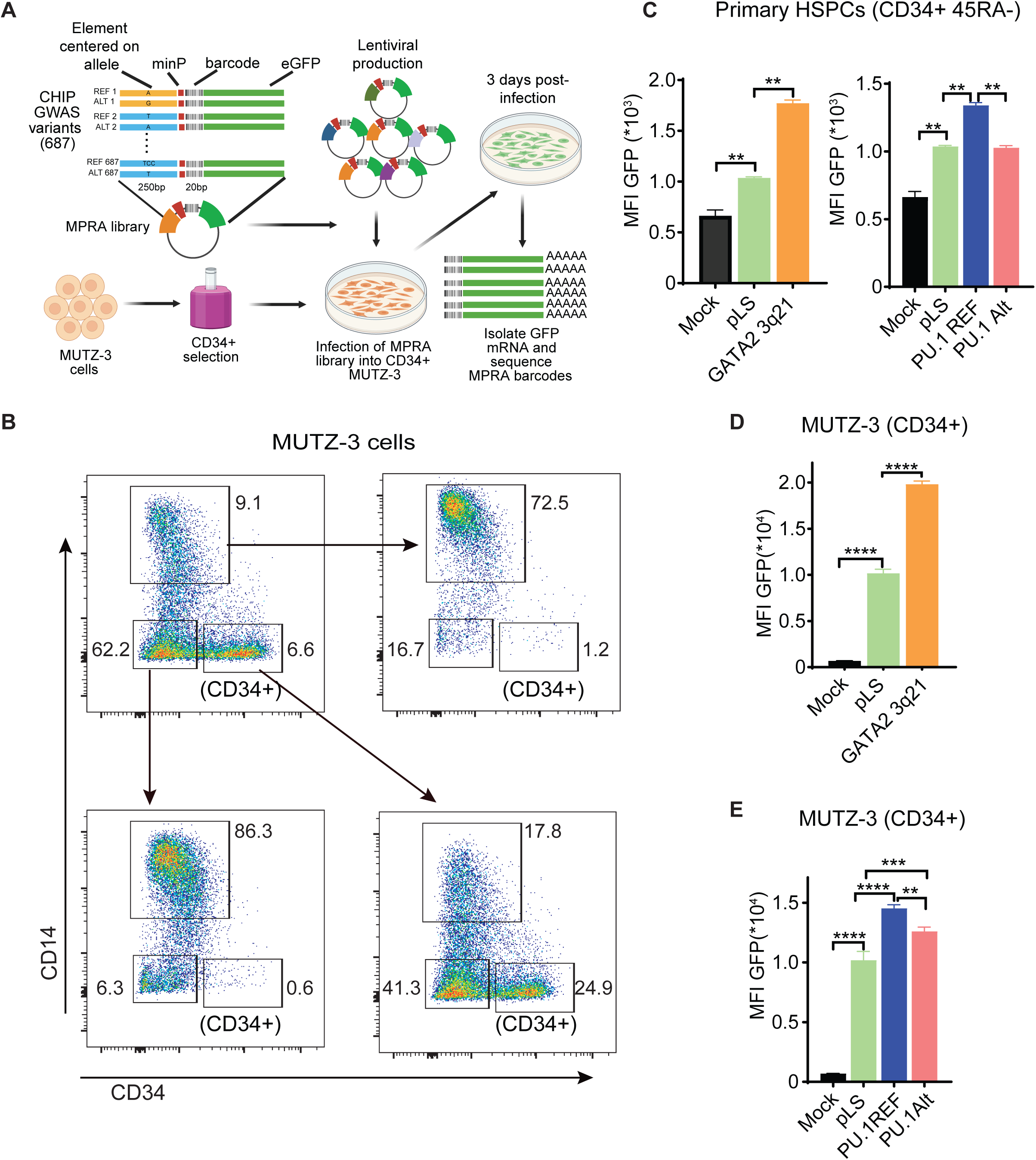
Human HSPC enhancers display similar reporter activities in primary human HSPCs and CD34+ self-renewing fraction of MUTZ-3 cells. (A) Schematic of the massively parallel reporter assay (MPRA) experiment to assess non-coding variants associated with CHIP. (B) FACS plots represent 3 distinct cell populations that were sorted from MUTZ-3 and cultured for 5 days. Results indicate that CD34+ population is the self-renewing population. (C) Reporter activity of PU.1 and GATA2 enhancers in primary human CD34+CD45RA-HSPCs as measured by GFP mean fluorescence intensity (MFI) using lentiviral reporter constructs (n=2 replicates). GATA2 3q21 indicate GATA2 oncogenic enhancer. PU.1 REF indicate PU.1 upstream regulatory element (URE) enhancer. PU.1 ALT indicate PU.1 URE with a mutation in a critical ETS motif within the enhancer. Empty reporter vector used in MPRA assay indicated by pLS. (D) GFP reporter activity of indicated enhancers in CD34+ fraction of MUTZ-3 cells 3 days post-infection (n=3 replicates). All data are means ± SDs. *p < 0.05, **p < 0.01 ***p < 0.001, ****p < 0.0001.

Since enhancers show cell-type specific activity, we searched for the appropriate cell-type to perform the MPRA. Since the disease relevant population in CHIP is the hematopoietic stem cells (HSCs), which is a rare population, we searched for alternative cell types. We selected MUTZ-3 cell line –a inv(3) AML cell line that is dependent on stem cell transcription factors such as MECOM and RUNX1^20,21^. MUTZ-3 cells also maintain a population of primitive CD34+ cells in culture that can self-renew or differentiate into CD14+ monocyte through an intermediate state **(Fig. 1B)**. Next, we confirmed if MUTZ-3 cells have trans-acting factors similar to primary HSCs, by testing two validated HSPC enhancers in lentiviral vectors with GFP reporter. We tested the GATA2 oncogenic enhancer that is commonly rearranged in inv(3) AML. We also tested the PU.1 upstream regulatory element (URE) that regulates PU.1 expression in HSCs. The empty MPRA vector without enhancers was sufficient to initiate GFP expression in primary CD34+ HSPCs and MUTZ-3 cells **(Fig. 1C, D)**. Both GATA2 and PU.1 enhancer further increased GFP expression compared to empty vector. Inactivating mutations in PU.1 enhancer reduced the GFP expression. The activity was similar to primary HSPCs in CD34+ population of MUTZ-3 cells **(Fig. 1C, D, E)**. Based on these results, we selected MUTZ-3 cells to perform the MPRA.

We infected the lentiviral MPRA library into 12 million CD34+ MUTZ-3 cells per replicate at an MOI of 33 providing an average of 242k integrations per sequence. 48 hours post-infection we lysed the MUTZ-3 cells for DNA and RNA and sequenced the barcodes by NextSeq. We confirmed that at this time point the MUTZ-3 cells still maintain the primitive CD34+ state (∼75%) and with no monocyte differentiation **(Fig. S1E)**. We were able to map the 96% of barcodes sequenced in MUTZ-3 cells back to enhancers using the plasmid enhancer-barcode associations. The experiments were highly reproducible, and all 5 replicates correlated well with each other at both the DNA and RNA barcode level **(Fig. 2A, B, Table S2)**. The positive control constructs behave as expected in the MPRA (PU.1 and GATA2 enhancers) **(Fig. 2C)**. In addition, we also included a CHIP risk variant containing TET2 enhancer from the TOPMed study which also showed allele specific activity **(Fig. 2C)**. Next, we examined the test constructs and found that they are highly represented with a median of 73 barcodes per replicates with high correlation between experimental replicates**(Fig. S1F, G, H, Table S2)**. We applied MPRAnalyze to estimate transcriptional activity (alpha) and observed constructs with significantly higher activity compared to the negative ORF controls **(Fig. 2D, Table S3)**^22^. We also find that activity of constructs that overlap with hematopoietic ATAC peaks was generally higher that non-ATAC constructs **(Fig. 2D)**. This shows that the assay is able to identify enhancers in an endogenous genomic setting. Having confirmed that the MPRA can detect endogenous HSPC enhancers, we next set out to identify constructs that showed significant allele specific activity based on CHIP variant (Ref vs. Alt allele) **(Table S4)**. We identified 87 variants that show significant differential allelic activity (qvalue < 10%; |log2 fold change| > 0.1) representing 12.9% of fine-mapped variants from 32 of 51 loci with a median of 1 MPRA hit/ GWAS loci **(Fig. 2E, Table S5)**. We were able to identify a single MPRA variant in 17 loci (**Fig. 2F)**. These results indicate that several CHIP risk variants may act on HSPC enhancers and can be prioritized by MPRA.

**Figure 2.**
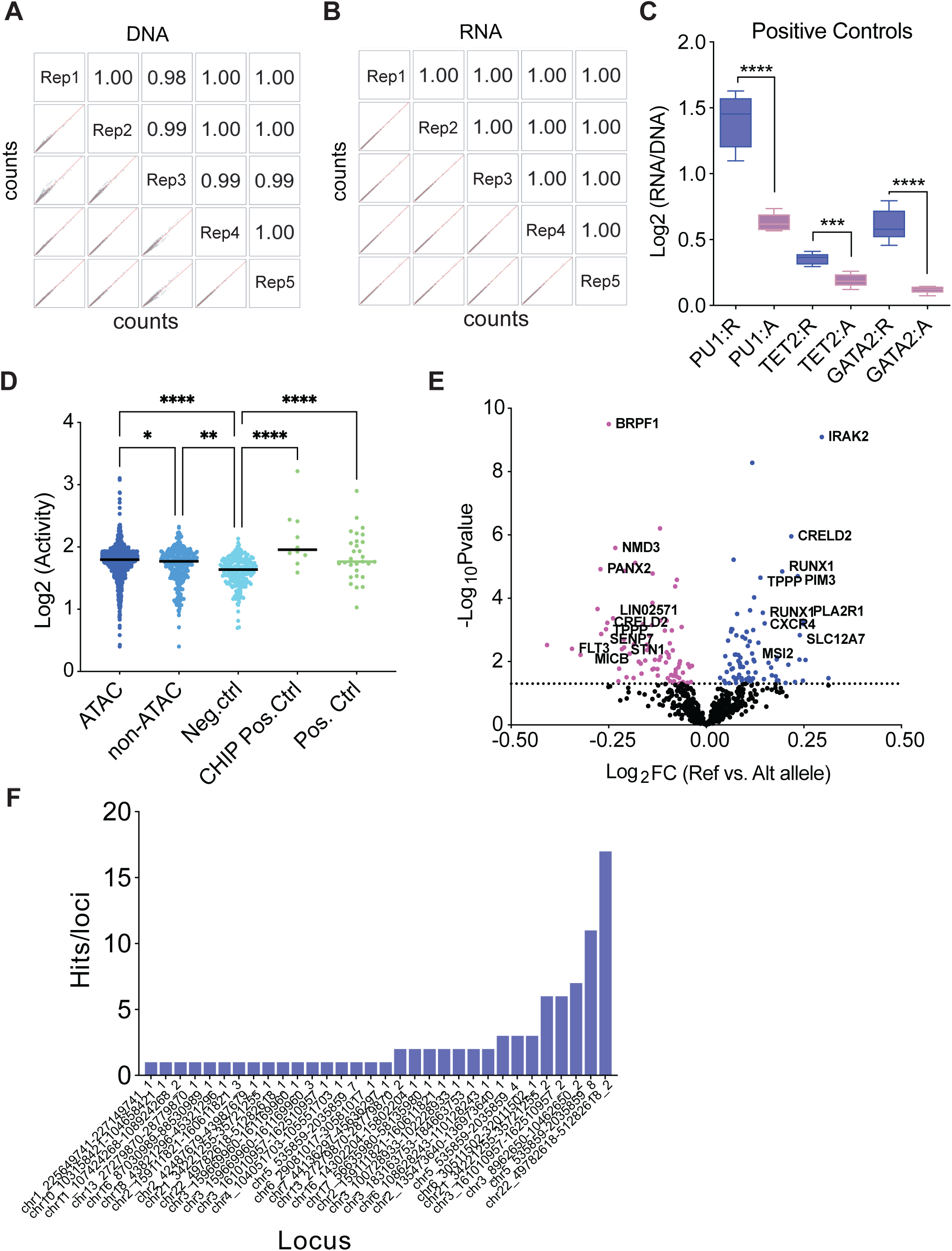
Identifying putative causal variants underlying CHIP predisposition loci using lentiviral MPRA in MUTZ-3 cells. (A) Correlation matrices of (A) DNA and (B) RNA barcode counts between 5 replicates. Correlation is provided as Pearson’s R coefficient. (C) Enhancer activity of positive control constructs reported as the log2-transformed RNA/DNA ratios calculated by summing RNA and DNA counts from all barcodes passing quality filters for each oligo. . (D) Enhancer activity (log2-transformed alpha) for oligos as reported by MPRAnalyze. Test constructs further grouped based on ATAC overlap. One-way ANOVA analysis with multiple comparisons performed. (E) Non-coding variants that show significant allele-specific activity (Ref vs. Alt) based on log2Fold change (FC) vs -log10 P-value. Nearest gene name is denoted for the hits. (F) Eighty-seven MPRA variants identified based on differential activity between reference (Ref) and alternate (Alt) alleles representing 32 GWAS loci (median = 1hit/locus). All data are means ± SDs. *p < 0.05, **p < 0.01, ****p < 0.0001.

Since MPRA is an exogenous assay, we next sought to validate endogenous activity of selected MPRA variants. We identified a MPRA variant rs35188965 that is located in the 1^st^ intron of the SLC12A7 gene. The variant is located in an open chromatin region in human HSPCs and disrupts a ETS motif, a well-known binding motif for several hematopoietic transcription factors such as PU.1, ERG, FLI1 and ETV6 **(Fig. 3A, S2A)**^23^. Using CRISPR/Cas9 we targeted the enhancer region containing the variant achieving high editing efficiency in hematopoietic cells **(Fig. 3B, S2C)**. Since SLC12A7 is not expressed in hematopoietic cells, we searched for gene expression changes in nearby genes post-editing and identified that NKD2^24^, a cell autonomous antagonist of the Wnt signaling pathway, is downregulated upon disruption of rs35188965 containing regulatory element, nominating NKD2 as a target gene affected by this variant **(Fig. 3C, S2B)**. We next targeted rs117145034, an MPRA variant located 150bp upstream of the FLT3 transcriptional start site and within open chromatin region in human HSPCs **(Fig. 3D, S2A)**. FLT3 encodes a receptor tyrosine kinase frequently mutated in AML^25^. Interestingly, rs117145034 variant modifies the spacing of tandem ETS motif creating closer binding sites potentially affecting cooperative binding **(Fig. 3D)**^26^. We disrupted the FLT3 variant containing region using CRISPR/Cas9 editing in the MOLM13 cells, an AML cell line that has surface expression of FLT3 and also contains a Flt3-ITD mutation **(Fig. S2D)**. Disruption of the variant harboring promoter region in MOLM13 cells resulted in downregulation of surface expression of FLT3 based on flow cytometry and decreased growth of these cells confirming the endogenous transcriptional activity of this region **(Fig. 3E, F, G)**.

**Figure 3.**
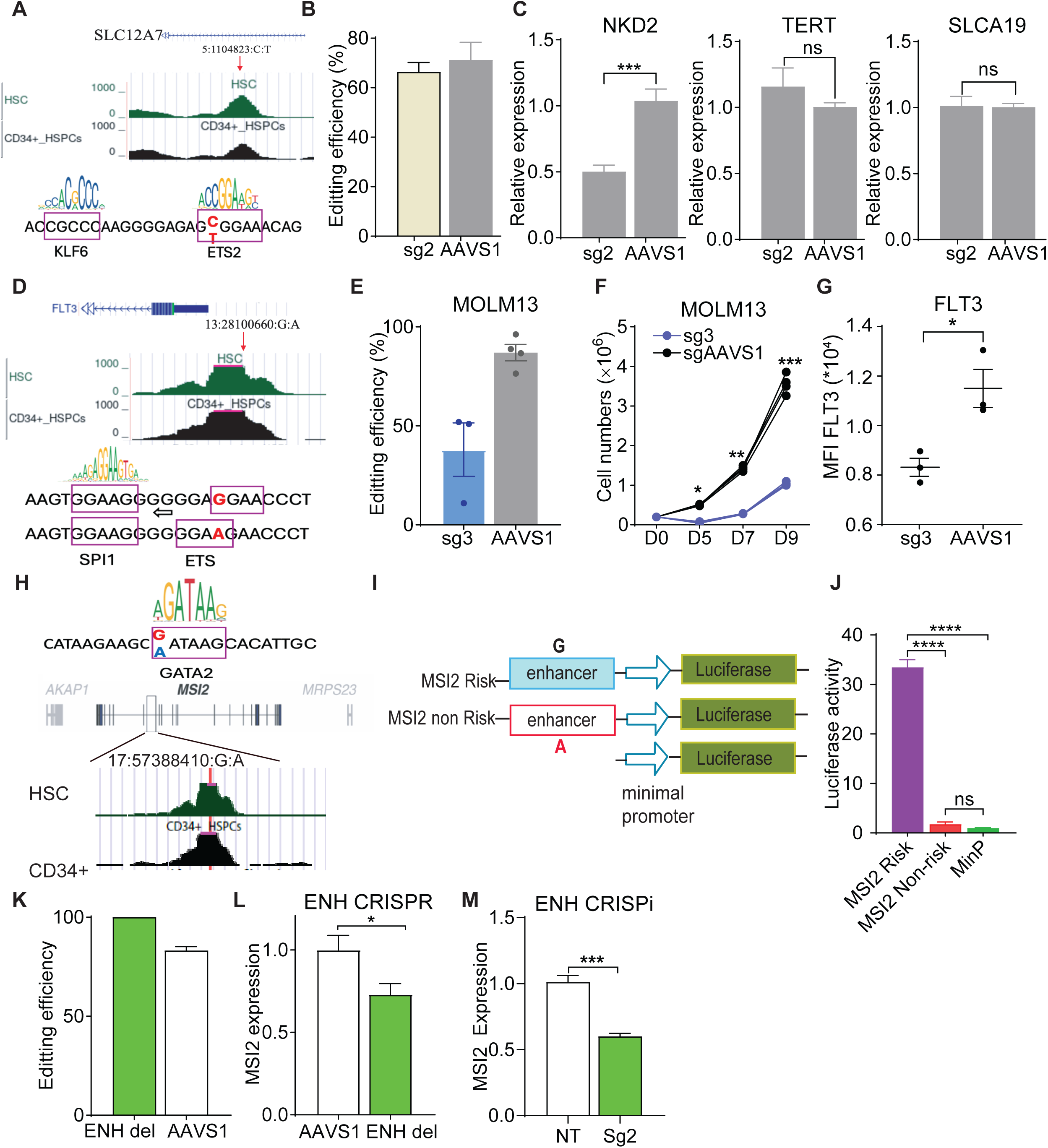
Validating endogenous activity of CHIP MPRA variants in hematopoietic cells using CRISPR genome and epigenome editing. (A) Genomic location of rs35188965 variant and accessibility in HSCs and motif analysis. (B) Bar graphs show relative expression of nearby rs35188965 genes by qPCR and CRISPR/Cas9 editing efficiency of the enhancer in K562 cells. (C) Genomic location of rs117145034 variant and accessibility in HSCs and motif analysis. (D) Growth curve of MOLM13 after electroporation with CRISPR/Cas9 sgRNAs. (E) FLT3 expression measured by FACS (n=3 replicates) and CRISPR/Cas9 editing efficiency at rs117145034 containing regulatory element in MOLM13 cells. (F) CHIP risk variant, rs17834140 located in 4th intron of MSI2 gene. (G) Schematic of luciferase constructs and luciferase reporter activity of Risk and non-risk alleles at rs17834140 (n=5). (H) Silencing of the variant containing enhancer using KRABdCas9 (n=3) and 51bp CRISPR deletion of the enhancer decreases MSI2 expression in K562s (n=3). (I) Frequency of edited alleles after CRISPR editing based on sanger sequencing analysis and deconvolution (TIDE) (n=3). All data are means ± SDs. *p < 0.05, **p < 0.01, ***p < 0.001, ****p < 0.0001.

To investigate how germline variants can modulate CHIP mutant clonal expansions we characterized rs17834140, a third candidate from our MPRA screen. This variant is located in an HSC accessible chromatin region within the 4th intron of the MSI2 gene **(Fig. 3H, S2A)**. We assessed the entire accessible chromatin region (∼500 bp) by reporter assay in hematopoietic cells and observed enhancer activity concordant with the MPRA screen with the protective “A” allele showing lower activity compared to the risk “G” allele likely by disrupting a GATA motif **(Fig. 3H, I, J)**. To confirm endogenous enhancer activity and identify target genes of the accessible region, we used CRISPR/Cas9 based microdeletions of this enhancer using 2 sgRNAs in hematopoietic cells **(Fig S2E)**. After three days post editing we observed a very high efficiency of microdeletions (∼90%) targeting rs17834140 and control AAVS1 edits (∼80%) **(Fig. 3K**). Edited cells showed a decrease in RNA expression of MSI2 upon disruption of the region **(Fig. 3L**). Independent epigenetic silencing using dCas9-KRAB confirmed this downregulation establishing MSI2 as the target gene of the enhancer **(Fig. 3M).**

Collectively, these results suggest the germline CHIP risk variant, rs17834140 modulates a hematopoietic enhancer with the risk “G” allele resulting in increased MSI2 expression and protective “A” allele preventing it. MSI2 is an RNA binding protein that regulates stability, splicing, translation and polyadenylation ^27^ and is a known driver of HSC self-renewal^28,29^. To determine if risk allele-mediated MSI2 upregulation affects clonal expansion in the context of CHIP, we overexpressed MSI2 using a lentiviral construct under control of an MSCV promoter to mimic the effect of the risk allele **(Fig. 4A)**. We overexpressed MSI2 in LSK cells from a well-established Tet2 knockout mouse model of clonal hematopoiesis and assessed self-renewal by CFU-C replating assays^30,31^ **(Fig. 4B)**. Consistent with previous studies, TET2 loss resulted in an increased self-renewal capacity of hematopoietic progenitors compared to wildtype (WT) controls as confirmed by colony forming activity until 3rd plating. However, MSI2 overexpression in Tet2-/- HSPCs resulted in a further increase in colony numbers and colony size compared to empty vector expressing Tet2-/- HSPCs **(Fig. 4C, Fig. S3C).** Similar increases in self-renewal were observed in Tet2+/- HSPCs upon MSI2 overexpression **(Fig. S3A-B)**.

**Figure 4.**
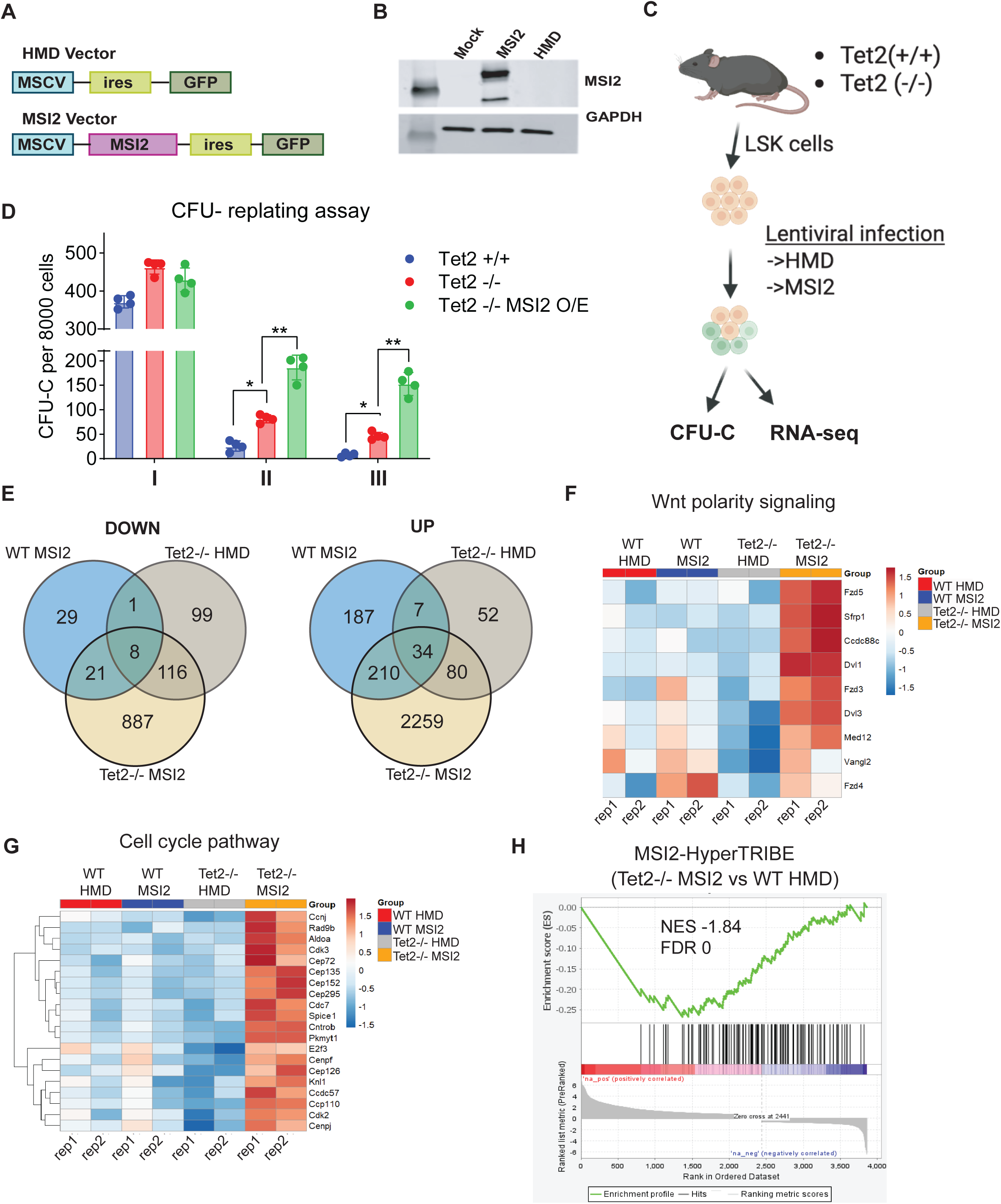
MSI2 enhances self-renewal of *Tet2* mutant murine HSPCs and is associated with upregulation of Wnt polarity signaling and cell-cycle pathways. (A) Schematic of lentiviral MSI2 overexpression constructs and Western blot confirming MSI2 overexpression in K562 cells. (B) Setup of MSI2 overexpression experiments in murine *Tet2*^-/-^ hematopoietic stem and progenitor cells (HSPCs). (C) Colony replating assays of sorted GFP^+^ Lin^-^Sca-1^+^c-Kit^+^ (LSK) cells shows increased self-renewal of *Tet2*^-/-^ HSPCs upon MSI2 upregulation. (D) Venn diagram showing common and unique differentially expressed genes across different genotypes identified by RNA-Seq (FDR <0.05 and |log2FC| >1.5). “WT MSI2” indicates wild-type HSPCs transduced with MSI2 overexpressing vector; “Tet2^-/-^ MSI2” indicates *Tet2*^-/-^ HSPCs transduced with MSI2 overexpressing vector; “Tet2^-/-^ HMD” indicates *Tet2*^-/-^ HSPCs transduced with empty HMD vector. All groups are compared against WT HSPCs transduced with empty HMD vector, “WT HMD” (n=2 replicates each group). (E) Heatmap showing relative expression of Wnt polarity genes across all groups. (F) Heatmap showing relative expression of Cell cycle genes across groups. (G) Gene set enrichment analysis of MSI2 HyperTRIBE binding targets in Tet2^-/-^ MSI2 vs. WT HMD. All data are means ± SDs. *p < 0.05, **p < 0.01.

To identify the pathways underlying the increased self-renewal, we performed RNA sequencing on Tet2-/- and WT LSK cells overexpressing MSI2 or control HMD vectors. Our analysis identified differentially expressed genes upon MSI2 overexpression in WT HSPCs (WT-MSI2 vs. WT-HMD; 2583 up and 1032 down genes) and upon MSI2 overexpression in Tet2-/- HSPCs (Tet2-/- MSI2 vs. Tet2 -/- HMD; 2442 up and 1417 down genes) **(Fig. S3F)**. We also identified differentially regulated genes upon just upon Tet2 loss (Tet2-/- HMD vs. WT-HMD; 173 up and 224 down genes) **(Fig. S3F)**. We observed that a majority of differentially regulated genes upon MSI2 overexpression in TET2-/- HSPCs did not overlap with either TET2 loss or MSI2 overexpression alone (887 downregulated and 2259 upregulated unique genes) **(Fig. 4E, S3D)**. Gene ontology (GO) analysis identified that Wnt cell polarity and cell cycle regulation pathways genes were upregulated upon MSI2 overexpression in Tet2-/- HSPCs **(Fig. 4F-G, S3G, Table S6)**. Among the top differentially upregulated genes in Tet2-/- HSPCs overexpressing MSI2 (KO-MSI2 vs. KO-HMD) were mitochondrial electron transport chain complex 1,3,4 genes (Nd1, Nd2, Nd4, Nd5, Nd6, Cytb, Co1, Rnr2) **(Fig. S3E, F)**. We also identified that the lncRNA MALAT1 which has been recently shown to be a downstream effector in TET2-/- hematopoietic cells was further upregulated upon MSI2 overexpression **(Fig. S3F)**^32^. Interestingly GO analysis identified that genes in the ATP production complex were downregulated suggesting MSI2 overexpression in Tet2-/- HSPCs may have diverse effects on mitochondrial metabolism **(Fig. S3G, Table S7)**. Previous studies have evaluated direct MSI2 binding targets in primary murine HSPCs using the HyperTRIBE approach overexpressing MSI2 fused to ADAR RNA-editing enzyme^33^. By performing GSEA analysis in Tet2-/- HSPCs overexpressing MSI2, using the murine HyperTRIBE dataset we identified an enrichment of MSI2 direct binding targets in murine HSPCs including several downregulated genes **(Fig. 4H)**. Collectively, our RNA-Seq results show that both MSI2 and TET2 gene networks are activated upon MSI2 upregulation in TET2-/- HSPCs collectively leading to increased clonal expansion. Furthermore, our results demonstrate a CHIP risk locus that modulates MSI2 expression in TET2 mutant HSPCs and thereby influencing self-renewal of TET2 mutant HSPCs.

## Discussion

While it is becoming increasingly clear that inherited genetic variation plays an important role in development of clonal hematopoiesis (CHIP), the underlying mechanisms driving this germline genetic risk still remains unclear^34,35^. A major challenge being that most of variants identified by GWAS on CHIP are located in the non-coding regions of the genome^36^. To overcome this, in our study we performed a lentiviral based MPRA on 687 non-coding variants from a GWAS on CHIP and identified 87 variants that show allele specific transcriptional activity in stem cell like populations. By complementing the MPRA results with CRISPR/Cas9 genome and epigenome editing approaches, transcription factor binding motif analysis and chromatin accessibility in hematopoietic stem cells we identified three variants that lie within endogenous enhancers and determined their target genes. Finally, we selected rs17834140 in the MSI2 locus for further functional studies and show that the CHIP risk allele mediated upregulation of MSI2 confers increased self-renewal capacity of TET2 mutant HSPCs. Our study highlights the utility of MPRA approach in localizing causal variants that underlie germline risk of clonal hematopoiesis.

MPRAs are powerful scalable approaches to identify non-coding causal variants associated with human diseases and traits. Previously we and others have utilized MPRAs to assess the transcriptional effects of non-coding variants and identify causal variants linked to red blood cell traits, autoimmune disorders, multiple myelomas, melanoma etc.^12,37,38^ Despite its successes the MPRA approach until recently was only feasible in diseases/traits where the affected cell types are amenable to episomal delivery approaches and yet to be applied in the context of CHIP. By using the lentiviral based integrating version of the MPRA we now evaluate CHIP risk variants for the first time in a stem cell relevant model system. Our lentiviral MPRA on CD34+ MUTZ-3 cells was able to reliably identify previously validated HSPC enhancers (PU.1 URE, GATA2 3q21, GATA2 9.5) and a validated CHIP risk variant (TET2 variant). Consistent with the enrichment of CHIP risk variants in accessible regions in HSPCs, the MPRA is now able to functionally identify several CHIP risk variants that influence transcriptional activity in stem cell populations, including the previously characterized African American specific CHIP risk variant at the TET2 locus. Similar to our previous studies on TET2 variant, three other putative variants identified from MPRA were validated. The MPRA identified a CHIP risk variant in the FLT3 locus that resulted decreased surface FLT3 expression. FLT3 is an FMS-like tyrosine kinase that is expressed in HSCs and is frequently mutated in AML^25^. Recent studies have shown that FLT3 is required for maintenance of leukemia stem cells and is dispensable for HSCs^39^. Our identification of FLT3 variant associated with CHIP risk, warrants further studies in examining the role of FLT3 expression in premalignant HSCs at the CHIP phase. We identified another variant in the intron of SLC12A7 that increases expression of the neighboring NKD2, a Wnt antagonist. Wnt signaling plays a critical role in self-renewal and maintenance of HSCs and overactive Wnt signaling leads to HSC exhaustion^40^. While aberrant Wnt signaling in BM microenvironment has been shown to lead to AML^41^, our results indicate that disruption of Wnt signaling may also play a cell-intrinsic role during CHIP development.

Finally, MPRA identified a variant in the intron of MSI2 gene that modulated MSI2 expression in hematopoietic stem cells. Our functional studies demonstrated that upregulation of MSI2 mimicking the risk allele conferred an increased *in vitro* self-renewal capacity to murine HSPCs harboring TET2 loss of function CHIP mutations. These results suggest a model by which germline genetic variants such as MSI2 interact with CHIP driver mutations to protect against clonal expansion of mutant HSCs. Our results are further supported by a recent work showing that loss of MSI2 can limit the clonal expansion of ASXL1 mutant human HSPCs^42^. We predict from CHIP driver specific GWAS analyses that the MSI2 variant may protect against several other CHIP driver mutations (such as DNMT3A and JAK2). How does upregulation of MSI2 cooperate with multiple CHIP driver mutations? Our RNA-seq analysis identified potential pathways such as mitochondrial metabolism, lncRNAs, cell cycle regulation and Wnt signaling pathways where such cooperativity can occur^32,43^. Our finding that MSI2 binding targets are enriched in TET2 KO/MSI2 overexpressing HSPCs, indicate that MSI2 networks are likely activated in CHIP mutant cells. Our findings also indicate the targeting MSI2 could be a potential strategy to prevent the development of CHIP mutant clones.

Our results indicate that CHIP risk variants can influence gene expression changes in HSPCs via a variety of mechanisms. For instance, the FLT3 and SLC12A7 variants both target the ETS motif. Instead of disputing the ETS motif, these variants either affect nearby sequences or affect spacing of a double motif. This can lead to disrupted binding of one of the ETS motif binding hematopoietic TFs, including FLI1, ERG, ETV4/6 and PU1. We also found that the MSI2 variant created a novel GATA2 binding motif likely leading to it upregulation. Taken together our results indicate that the CHIP risk variants likely affect the binding of the heptad transcription network (FLI1, ERG, GATA2, RUNX1, TAL1, LYL1, and LMO2) that is critical for regulation of expression of HSPC self-renewal and maintenance genes^44^.

We recognize that MPRA approach is subject to certain limitations. MPRA variants were not identified for 37% (19/51) of CHIP GWAS loci suggesting that causal variants at these loci may have alternative mechanisms. For example, causal variants at these loci can function through alternative splicing or by modulating transcript stability. Causal variants can act via cell-extrinsic mechanisms by targeting the BM microenvironment to increase CHIP risk. Another potential issue is that most common CHIP driver mutations affect epigenetic factors (TET2, DNMT3A, ASXL1) or splicing factors (SF3B1, SRSF2) that can also influence the effects of germline variants on cis-regulatory elements or completely change the regulatory landscape in CHIP mutant cells. In order to address this future MPRA studies (assessing mRNA expression, splicing and stability) can be implemented into primary HSPCs or iPSC derived HSPCs engineered to carry CHIP mutations and/or cell types within the BM microenvironment.

In summary, the MPRA approach provides a catalog of genetic variants associated with CHIP that affect transcriptional regulatory elements in HSPCs. Intersecting MPRA results with orthogonal approaches such as statistical fine mapping, regulatory element mapping, gene-centric loss of function screens in CHIP relevant cell types will enable us to rapidly identify causal mechanisms underlying CHIP predisposition.

## Acknowledgments

We are grateful for the members of the Nandakumar laboratory for valuable comments and suggestions. This work was supported by the Vera and Joseph Dresner Foundation MDS Research Fund (S.N) and Edward P. Evans MDS Foundation Discovery Research Grants (S.N) and the startup funds from the Albert Einstein College of Medicine (AECOM) and the Montefiore Einstein Comprehensive Cancer Center (MECCC). This work was also supported by US National Institutes of Health grant R35HG011329 (R.T).

## Author Contributions

T.N., J.J., R.T., and S.N. conceptualized the study and devised the methodology. T.N., J.J., S.K., D.K., A.H., A.S performed experiments. J.C performed computational analysis. C.Y., K.I., R.V., and K.S provided resources and feedback. T.N., J.J., and S.N. wrote the original manuscript, as well as edited the manuscript with input from all authors. S.N acquired funding for this work and provided overall project oversight.

## Supplemental Tables Legends

Table S1: Constructs Synthesized for MPRA Targeting CHIP GWAS Variants

Table S2: Normalized RNA and DNA barcode counts for all 5 MPRA replicates from MPRASuite pipeline

Table S3: Enhancer activity for 1588 constructs captured in MPRA using MPRAnalyze

Table S4: Allelic skew measurements of 672 CHIP GWAS associated variants using MPRAnalyze

Table S5: Eighty seven MPRA hits and their annotations

Table S6: Gene ontology analysis of unique upregulated pathways - Tet2^-/-^ MSI2 vs. WT HMD

Table S7: Gene ontology analysis of unique downregulated pathways - Tet2^-/-^ MSI2 vs. WT HMD

Table S8: List of primers used for MPRA library construction and processing

Table S9: List of primers used for RT qPCR and PCR assays

Table S10: List of sgRNAs used for CRISPR and CRISPRi experiments

## Methods

### Individual lentiviral reporter assays in primary human HSPCs

The lentiviral reporter constructs were designed to deliver known human HSPC enhancer elements containing the GATA2 3q21 and PU.1 URE enhancers that were positioned upstream of a minimal promoter and driving the reporter GFP. These constructs were generated using the LentiMPRA backbone (pLS-SceI). Fragments containing a 300bp human GATA2 3q21 enhancer (hg38, chr3:128,603,821-128,604,120) and 200bp PU.1 URE enhancer element (hg38, chr11:47,394,824-47,395,023) were synthesized as gBlocks (IDT technologies) together with the adapter sequence (15bp) and minimal promoter (32bp) used in the original LentiMPRA design and cloned using SbfI and AgeI sites into the pLS-SceI backbone. To generate the mutant PU.1 URE (Alt) a critical ETS binding site “GGAA” at position 135-138 was mutated to “TCGC” similar to a previous study^45^. Lentiviral supernatants of these constructs were produced by transient transfection in 293Ts and were used for subsequent transduction of human HSPCs. Primary CD34+ human HSPCs from adult mobilized peripheral blood were thawed 48hrs before transduction and cultured in human HSC maintenance conditions with StemSpan SFEM II media with 1x CC100 cytokine cocktail (StemCell Technologies) and human TPO (50ng/ul) and small molecule UM729 (500nM). The HSPCs were transduced with the lentiviral reporters by spinfection at 2400rpm for 90 min in the presence of 8 µg/mL Polybrene (Millipore). The cells were cultured for 5 days post-transduction in HSC maintenance conditions and reporter GFP expression in primitive HSPC compartment was measured by flow cytometry. The panel of antibodies to identify human HSPC compartment were anti-CD45RA-APC-H7 (BD, 560674) and anti-human-CD34-BV421 (Biolegend, 343610).

### Individual Ientiviral reporter assays in MUTZ-3 cells

MUTZ-3 cells (DSMZ) were cultured at 37 °C in α-MEM (Life Technologies) supplemented with 20% FBS, 20% conditioned medium from 5637 cells (ATCC) and 1% penicillin/streptomycin. Confluency was maintained between 5 × 10^5^ and 1.5 × 10^6^ per ml. 5e5 MUTZ-3 cells were transduced with lentiviral supernatants of reporter constructs containing GATA2 3q21 and PU.1 URE enhancers by spinfection at 2400rpm for 90 min in the presence of 8 µg/mL Polybrene (Millipore). The cells were cultured for 3 days post-transduction and reporter GFP expression in primitive CD34+ compartment was measured by A5 Symphony flow cytometer. The panel of antibodies used to identify primitive CD34+ compartment of MUTZ-3 cells were CD34-APC (BioLegend, 343608), CD14-PE-Cy7 (BioLegend, 367112). For experiments to identify the primitive compartment of MUTZ-3 cells, FACS sorting for 3 distinct subpopulations of MUTZ-3 cells were performed in BD Aria FACS sorter. The three sorted populations were cultured in MUTZ-3 media for 5 days and FACS analysis was performed again using CD34-APC (BioLegend, 343608) and CD14-PE-Cy7 (BioLegend, 367112).

### Variant selection for MPRA

Variant selection for MPRA: A total of 3,842 CHIP associated SNPs across 51 loci identified using Bayesian fine-mapping approach (95% credible set) were obtained from Supplementary Table S4 in Kessler et al study (Kessler et al., Nature 2022). The ATAC-seq peaks across 18 primary human hematopoietic populations (29August2017_EJCsamples_allReads_500bp.bed) was downloaded from Ulirsch et al study (Ulirsch et al Nature Genetics 2019). After converting hemeATAC peaks from hg19 to hg38 using the UCSC LiftOver tool the ATAC peaks were intersected with 3,842 CHIP associated variants using bedtools with the following options “bedtools intersect -a atac_file_38_500.bed -b 3842_gwas.bed -wa -wb “. This analysis yielded 560 CHIP associated SNPs spanning 39 loci that overlapped with hemeATAC peaks. In addition, we selected SNPs with high posterior inclusion probability (PIP) scores (0.05–1) that were located outside ATAC-seq peaks, resulting in 127 additional SNPs across 41 loci. Considering both the reference and alternate alleles of the overlapping and non-overlapping SNPs, we generated a set of 1,374 test variants. For positive controls we included validated HSPC enhancers such as GATA2 oncogenic enhancer, GATA2 +9.5 enhancer variant (rs1559985787) identified in MonoMac syndrome, PU.1 URE and a TET2 CHIP risk variant (rs144418061) identified in our previous work (10 HSPC positive controls). Our library also contained an additional 30 previously validated positive controls from multiple studies. For negative controls, 200 open reading frames (ORFs) corresponding to coding variants were incorporated. In total, these selections produced a library of 1,614 enhancer elements. Oligos were synthesized (Twist Biosciences) as 280 bp sequences containing 250bp of genomic sequence centered around the variant (124 bp on the 5’ and 125 bp on the 3’ end for SNPs) and 15 bp of adaptor sequence on either end **(Table S1)**. For oligos with insertion/deletions, the 5’ and 3’ flanking genomic sequences were proportionately trimmed with respect to the size of the indel. These oligos have the same start/stops with respect to the reference coordinates of the oligo sequence (i.e. total sequence length was discordant between “Ref” and “Alt” due to the presence or absence of the indel). Total oligo length of the library never exceeded 280bp including flanking adapters.

### MPRA library construction

The MPRA libraries were constructed as previously described with several modifications to the original protocol^15^. Sequences were synthesized as 280 bp oligonucleotides libraries by Twist Biosciences. Unique 20 bp barcodes were added by PCR using primers #82 and #202. Each oligo amplification consisted of 24 50 uL reactions containing 25ul Q5 Ultra II Master Mix, 2.5uL each of 10uM primer, and 0.5 uL oligo library using the following PCR conditions (98°C for 30s; 8x (98°C for 10s; 60°C for 15s; 65°C for 45s); 72°C for 5m). PCR products were purified using AMPure XP SPRI (Beckman Coulter, A63881) and incorporated into the SfiI digested pLS-SceI:MPRAv3-MCS backbone vector by Gibson assembly (35uL 2xNEB HiFi Assembly MM, 30ngg DNA oligo pool, 90ng SfiI digested pLS-SceI:MPRAv3-MCS in 70uL reaction incubated at 50°C for 1 hour). To achieve a library composition of ∼200 barcodes per oligo, a test transformation was performed with 1uL of the 20uL SPRI purified Gibson assembly mixture and 50uL of Endura Electrocompetent Cells (Biosearch Technologies, 60242) to determine the optimal amount of Gibson assembly and competent cells to achieve the desired colony-forming unit (CFU) count. Based on the results of the test transformation, a subset of a second identical transformation was taken directly after electroporation and split across ten 1 mL cultures with recovery media then incubated for 1 hour at 37°C. After 1 hour, each culture was independently expanded in 20 mL of Terrific Broth (TB) with 100ug/mL of carbenicillin. In parallel, CFU colony counting plates were created from 4 of the 10 cultures. After 18 hours of growth at 30°C, the cultures were individually pelleted and frozen, and the desired number of cultures from 10 expanded cultures was selected, based on the CFU counting plates, to reach an average of ∼200 CFUs (barcodes) per oligonucleotide. Cultures were combined and purified using a Qiagen Plasmid Plus Maxi, followed by further purification by Monarch PCR & DNA Cleanup Kit (NEB, T1030) . Fifteen colonies from the CFU plates were checked by colony PCR to determine the oligo insertion rate for the test transformation with all plasmids containing an oligo insert. The expanded purified pLS-SceI:MPRAv3-MCSΔminP plasmid library was sequenced using Illumina 2x150 bp chemistry to acquire oligo-barcode pairings. To construct the final MPRA libraries, 10ug of the pLS-SceI:MPRAv3-MCS:ΔminP library was digested with AsiSI, and a minP promoter (amplified from pMPRAv3:minP-GFP, Addgene #109035) was inserted using Gibson assembly (75uL 2xNEB HiFi Assembly MM, 250ng minP amplicon, 1ug pLS-SceI:MPRAv3-MCS:ΔminP in a 150uL reaction incubated at 50°C for 1.5 hours). The Gibson reaction was purified using 1x SPRI and eluted in 40 uL water prior to electroporation of 8 uL of purified plasmid library into 200 uL Endura cells. The electroporation was split across six 2 mL cultures and recovered at 37°C for one hour followed by expansion of each 2 mL culture in 500 mL of TB media with 100ug/mL of carbenicillin. After 16 hours of growth at 30°C, plasmid was purified using Nucleobond RS column. CFU counting plates from 4 of the 500 mL cultures were made to monitor transformation efficiencies (50 million). Thirteen colonies from the CFU plates were checked by colony PCR to determine the minP insertion rate for the test transformation with 12 of 13 plasmids containing a minP insert.

### MPRA library viral production and titration

Lentivirus was produced by transfecting 293T cells with the viral packaging plasmids pVSV-G and pΔ8.9, along with the MPRA plasmid library, using calcium phosphate transfection method. After 24 hours, the transfection medium was replaced with viral collection media containing DMEM (Life technologies) supplemented with 10%FBS, 10mg/ml BSA and 1% penicillin/streptomycin. Viral supernatants were collected 24 hours later, filtered through a 0.45 µm filter, concentrated by ultracentrifugation at, 22,000rpm for 90 minutes at 4C and resuspended in MUTZ-3 media and stored at –80 °C. For viral titration, frozen concentrated virus stocks were thawed, and varying volumes were used to infect 4 × 10⁵ MUTZ-3 cells in the presence of Polybrene (8 µg/mL). Infections were carried out by Spinfection at 2000 rpm for 90 min at 22 °C. After 42 h post-infection, GFP expression was analyzed by flow cytometry and genomic DNA extracted and the multiplicity of infection (MOI) was measured by quantitative PCR (qPCR) using primers that amplify amplify viral DNA (WPRE.F and WPRE.R). genomic DNA (LP34.F and LP34.R) and plasmid backbone (BB.F and BB.R) **(Table S9).**

### Setup of the MPRA experiment

The CD34+ primitive population of MUTZ-3 cells was isolated using CD34 magnetic bead separation (EasySep, StemCell Technologies) according to the manufacturer’s instructions. The purity of the isolated cells was confirmed by flow cytometry following staining with CD34-APC (BioLegend, 343608), CD14-PE-Cy7(BioLegend, 367112) antibodies. A total of 1.2 × 10⁷ CD34+ MUTZ-3 cells were transduced with the concentrated lentiviral MPRA library stocks at a multiplicity of infection (MOI) of 33. Transductions were performed in six independent replicates with polybrene (8 µg/mL). Spinfection was carried out at 2000 rpm for 90 min at 22 °C, after which the cells were incubated overnight in viral supernatant. After 17 hours, the culture medium was replaced with fresh MUTZ-3 medium. At 48 h post-transduction, cells were harvested for RNA and DNA isolation, and a small subset was collected for flow cytometry (FACS) for GFP expression and multiplicity of infection (MOI) quantitative PCR (qPCR) as described above.

### RNA isolation and MPRA RNA library generation

Total RNA was extracted from the cell homogenates using the Quick-DNA/RNA MagBeadi GFP mRNA was then pulled down using a mixture of 3 GFP-specific biotinylated primers at 0.5 nM (#843, #123 and #844, **Table S8**) in 0.2x SSC and 33% Formamide (MilliporeSigma 4650-500ML) and incubated for 2.5 hours at 65°C. Biotin probes hybridized with GFP mRNA were then captured by adding 400 uL of pre-washed Sera Mag Beads (Fisher Scientific) eluted in 500 uL of 20X SSX by agitation at room temperature for 15 minutes. The beads were captured on a magnet and washed once with 1X SSX and twice with 0.1X SSC. After removal of the last wash, 50 uL of water was added and the RNA was treated 1 hour with Turbo DNase at 37°C. The beads were then collected by magnet and the supernatant removed and purified with 2x RNA SPRI. Complementary DNA was created using SuperScript III (Life Technologies) in a 100 uL reaction with a gene specific primer for the GFP transcript (20 uM primer #851, **Table S8**) and a modified elongation temperature (47°C for 80 minutes). GFP mRNA abundance was quantified by qPCR to determine the cycle threshold for each replicate using a QuantStudio 5 qPCR instrument (Reaction mix: 5 uL Q5 Ultra II 2x, 0.5 uL 10uM primers #845 and #802, 1.66 uL 1:10,000 Sybrgreen 1, 1 uL cDNA or GFP mRNA in a 10 uL reaction, Cycle conditions: 98°C for 30s; 40x [98°C for 10s; 62°C for 15s; 72°C for 30s], **Table S8**). A standard curve with the final MPRA library (10 fg - 1 ng) was run alongside 1 uL of cDNA and 1 uL of GFP mRNA for each replicate. Any samples showing amplification in the mRNA sample were discarded for having plasmid carryover. cDNA replicates within a cell-type were diluted to approximately the same concentration based on the qPCR results, and first round PCR (98°C for 30s; 4x [98°C for 10s; 62°C for 15s; 72°C for 30s]; 72°C for 2m) with primers #845 and #802 **(Table S8)** were used to amplify barcodes associated with GFP cDNA sequences for each replicate. A second round of PCR (98°C for 30s; 14x [98°C for 10s; 68°C for 15s; 72°C for 30s]; 72°C for 2m) was used to add Illumina sequencing adaptors. Genomic DNA replicates within a cell type used 200ng per reaction and 8 reactions with the first round PCR (98°C for 30s; 46-13x [98°C for 10s; 62°C for 15s; 72°C for 30s]; 72°C for 2m) with primers #845 and #802 **(Table S8).** Next in strip tubes reverse SPRI was performed at 0.5X and then 1.7X to eliminate genomic DNA. Serial elute with 30 uL of EB. The second round of PCR (98°C for 30s; 19x [98°C for 10s; 68°C for 15s; 72°C for 30s]; 72°C for 2m) was used to add Illumina sequencing adaptors. The resulting MPRA barcode libraries were spiked with 0.01-1% PhiX and sequenced on an Illumina NextSeq 500 or NovaSeq SP.

### MPRA data processing and main analysis

MPRA data were analyzed as previously described^15^ using the MPRASuite pipeline (https://github.com/tewhey-lab/MPRASuite), comprising MPRAmatch (barcode/oligo pairing) and MPRAcount (tag assignment and counting). Briefly, oligo-barcode pairs were identified by merging paired-end reads from the lenti transfer plasmid into single amplicons using Flash2 (2.2.00)^46^. Genomic DNA sequences were extracted and mapped to the oligo design file using minimap2 (v2.17-r941)^47^. To ensure high-quality pairing, alignments with greater than 5% error were discarded, and valid pairs were compiled into a lookup table. Following barcode tag sequencing of RNA and genomic DNA libraries, 20-bp barcode tags were assigned to oligos using this lookup table. Count data was filtered so barcodes that did not have counts of in both DNA and RNA for a replicate were removed for that replicate. Oligos with 2 or fewer paired barcodes in each of the replicates were further removed. Quantitative analysis was performed using the MPRAnalyze ^22^. Sequencing depth factors were estimated via the upper quartile method, correcting for library size differences across batches. Transcriptional activity (alpha) was modeled with DNA counts controlled for batch effects (∼batch) and RNA counts modeled using an intercept-only design (∼1). Empirical p-values were calculated using ORF negative controls to establish the null distribution. To test for allelic effects, both alleles were analyzed using MPRAnalyze’s comparative mode where DNA counts were modeled with replicate, barcode, and allele effects (∼replicate + barcode + allele), and RNA counts were modeled with replicate and allele effects (∼replicate + allele). Allelic skew was assessed using a likelihood ratio test comparing the full model to a reduced model without the allele term (∼replicate). Multiple testing correction was performed using Storey’s q-value method.

### Enhancer disruption and deletion using CRISPR/Cas9

SpCas9 compatible **s**ingle or double sgRNAs to either disrupt or delete MPRA variant containing enhancers were designed using the Benchling platform (www.benchling.com) and then ordered as modified sgRNA (Synthego) **(Table S10)**. Briefly, the RNP complex was made by combining 100 pmol Cas9 (IDT) and 100 pmol modified sgRNAs (Synthego) and incubating at for 15 min at room temperature. For samples that underwent deletion with 2 guides, total amounts of 100 pmol Cas9 and 100 mol sgRNA (50 pmol each guide) were used. Either 2 × 10^5^ K562 cells or MOLM13 cells were resuspended in 20 µl SF solution were mixed with RNP complex and underwent nucleofection with program FF-120 (K562s) or EZ-100 (MUTZ-3) in Lonza 4D nucleofector instrument with 20 µl Nucleocuvette strips. At day 3 or day 5 post-electroporation, DNA and RNA were extracted from edited cells using the AllPrep DNA/RNA Micro kit (Qiagen) according to the manufacturer’s instructions. Genomic PCR was performed using Platinum II Hotstart Mastermix (Thermo Fisher Scientific) and edited allele frequency was detected by Sanger sequencing and analyzed by ICE (ice.syngthego.com). The effect on putative target gene mRNA expression after editing was detected by quantitative PCR with reverse transcription (qRT–PCR) using SYBR green (ThermoFisher scientfic) after cDNA synthesis with iScript (Bio-Rad) with specific primers **(Table S9)**. The effect on FLT3 surface expression in MOLM13 cells post-editing was measured by flow cytometry using anti-human CD135 (FLT3, BD ebioscience, 567273) antibody by A5 Symphony flow cytometer. The MOLM13 and K562 cells used in these experiments were cultured in RPMI media supplemented with 10% FBS, 1% penicillin/streptomycin.

### Enhancer silencing using CRISPR interference

For CRISPRi experiments we used an all-in-one lentiviral vector backbone expressing sgRNA driven by a U6 promoter, and MSCV promoter driving expression of a KRAB domain fused to dCas9 linked to GFP reporter (by a self-cleaving 2A peptide sequence from porcine teschovirus). The sgRNAs targeting MSI2 enhancer and non-targeting control were cloned into this vector using BsmBI restriction enzyme sites similar to previous studies^48^ **(Table S10).** K562 cells were infected as described above and GFP+ cells were FACS sorted 48 h post-infection and expanded for an additional 3 days. RNA was extracted using a RNeasy Mini kit (QIAGEN) and MSI2 expression was tested by quantitative PCR as described above.

### Luciferase reporter assays

The genomic region containing risk and non-risk alleles of the variant rs17834140 (∼500bp) were synthesized as gBlocks (IDT Technologies) and cloned into the Firefly luciferase reporter constructs (pGL4.24) using KpnI and EcoRV sites. Each allele-specific construct differed by only the single nucleotide of the variant of interest (i.e., all other nucleotides in the construct are the same). The Firefly constructs (500ng) were co-transfected with pRL-SV40 Renilla luciferase constructs (50ng) into 100,000 K562 cells using Lipofectamine LTX (Invitrogen) according to manufacturer’s protocols. Cells were harvested after 48 hours and the luciferase activity measured by Dual-Glo Luciferase Assay system (Promega). For each sample, the ratio of firefly to Renilla luminescence was measured and normalized to the minimal promoter construct. Location of rs17834140 gblock in hg38 is chr17:57,388,064-57,388,601.

### Generation of MSI2 overexpressing lentiviral constructs

To generate the MSI2 overexpression construct the MSI2 cDNA sequence from GENCODE v43 was codon optimized and synthesized as gBlocks (IDT technologies) together with flanking EcoRI and XhoI restriction enzyme sites and cloned into the HMD lentiviral backbone linked to GFP reporter (by ires sequence). Concentrated lentiviral particles were produced as described above for lentiMPRA library.

### Mouse models

Tet2^-/**-**^ mice were previously characterized in the laboratories of Drs. Keisuke Ito and Meelad Dawlaty^30^. This is a well-established model of clonal hematopoiesis characterized by expansion of HSPC compartment before development of myeloid malignancies and increased hematopoietic repopulating ability. The mice were established and housed in an SPF (Specific-pathogen-free) barrier facility. Animals with no overt signs of hematologic disease based on peripheral blood analyses were sacrificed at 8-12 weeks of age to harvest bone marrow for colony forming (CFU-C) assays and RNA-seq analyses. All studies were conducted in accordance with our IACUC approved protocols overseen by the Institute for Animal Studies at the Albert Einstein College of Medicine.

### CFU-C colony replating assays

Bone marrow cells from 8-12 weeks old TET2^-/-^ mice, TET2^+/-^and littermate wildtype (WT) were flushed in sterilized DPBS, then were enriched for hematopoietic stem and progenitor cells using EasySep^TM^ mouse Hematopoietic Stem and Progenitor Isolation kit (Stem Cell Technologies). TET2^-/-^ and WT LSK cells were sorted and cultured overnight in STEMSPAN (Stem Cell Technologies) containing 1% L-Glutamine, 1% P/S, murine SCF (50ng/ml), murine TPO (50ng/ml) at 37 °C. The following day, 50,000 TET2^-/-^ LSK cells were spinfected with concentrated HMD and MSI2 lentiviral particles at 2000rpm, 22 degrees Celsius, for 90 minutes. They were then incubated for 20 hours and subsequently changed media. Two days later, GFP+ cells were sorted and plated at 5000 cells per plate using Mouse Methylcellulose Complete Media HSC007 (R&Dsystems). Every 7 days colonies were counted and imaging performed using EVOS™ M7000 Imaging system (ThermoFisher Scientific). Post-imaging the colonies were disrupted into single cell suspensions and 8000 cells were replated for subsequent CFU-C plating. A total of 3 colony replating was performed.

### Bulk RNA Seqeuncing and analysis

TET2^-/-^ and WT LSK cells were infected with HMD and MSI2 lentiviral vectors as described above and FACS sorted GFP+ cells 2 days post-infection were lysed for RNA using RNeasy Plus Micro Kit (Qiagen), and bulk RNA seqeuncing was performed using Poly-A enrichment method (2 biological replicates per group) (Admera Health, NJ). Library preparation were carried out with the NEBNext Ultra II Directional RNA Library Prep Kit and sequencing was performed on the Illumina NovaSeq 6000 platform, generating approximately 40 million reads per sample (Admera Health, NJ). Bulk RNA-seq was performed in two biological replicates using primary HSPCs (LSK cells) isolated from a well-established TET2 mouse model. Total RNA was extracted using the RNeasy Mini Kit, and poly(A) enrichment and library preparation were carried out with the NEBNext Ultra II Directional RNA Library Prep Kit. Sequencing was performed on the Illumina NovaSeq 6000 platform, generating approximately 40 million reads per sample. Raw read quality was assessed using FastQC v0.11.8, and adaptor trimming and removal of low-quality bases were performed with Trimmomatic v0.38^49^ using default settings. Reads were aligned to the mm39 genome using the STAR v2.7.1a aligner^50^, and alignment files were processed with Samtools v1.9 to filter low-quality reads^51^. Gene-level quantification was performed with HTSeq^52^, and differential expression analysis was conducted using DESeq2^53^. Gene set enrichment analysis was carried out using GSEA v4.3.2^54^, incorporating the HyperTRIBE-identified MSI2 target gene list^33^. Gene Ontology (GO) enrichment analysis was performed with the DAVID Functional Annotation Tool^55^.

---

**Supplemental figure 1:**
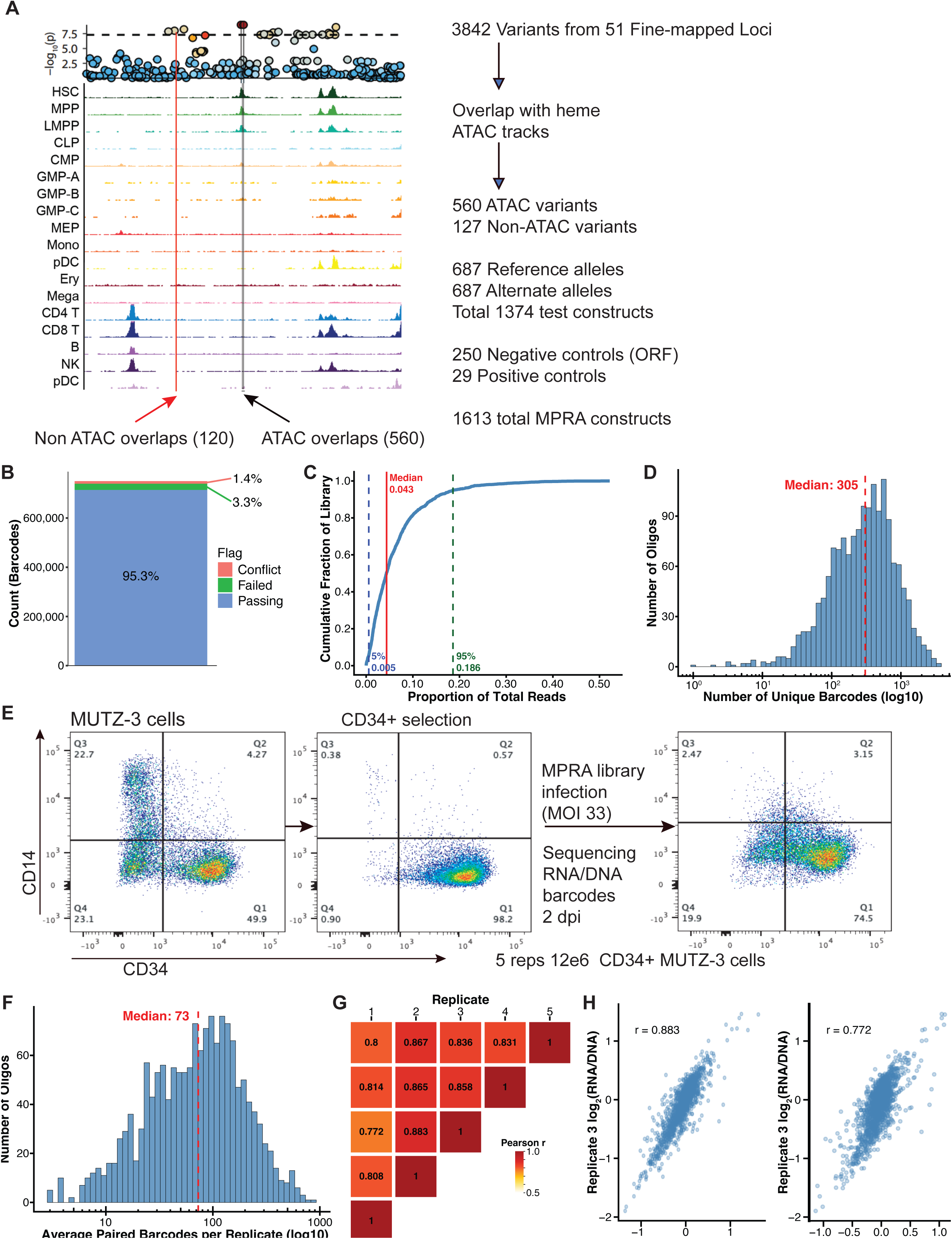
(A) Representative Locus zoom plot showing association probabilities and chromatin accessibility across a various stage of hematopoiesis and the strategy used for selecting variants to be tested in MPRA. (B) Percentage of barcode-oligo associations in the MPRA library that passed sequencing and mapping quality filters. (C) Cumulative distribution function showing MPRA library uniformity. The x-axis represents the proportion of total reads across all barcodes in the plasmid library for each oligo. Dashed lines indicate the read proportions at the 5th and 95th percentiles. Perfect uniformity would yield a proportion of 0.062 (1/1614) for each oligo. (D) Number of distinct barcodes associated with each oligo in the lenti transfer plasmid library. (E) Schematic of lentiviral library transduction into sorted CD34+ cells for MPRA screening. (F) Histogram of the median number of paired barcodes per oligo across all five MUTZ-3 replicates. Only barcodes with counts in both DNA and RNA that passed all quality filters are included. (G) Correlation matrix showing Pearson’s r for log2-transformed RNA/DNA ratios between individual MUTZ-3 replicates. Ratios were calculated by summing RNA and DNA counts from all barcodes passing quality filters for each oligo. (H) Representative scatter plots of log2-transformed RNA/DNA ratios comparing individual MUTZ-3 replicates. Replicate pairs with the lowest and highest Pearson’s r values are shown.

**Supplemental figure 2:**
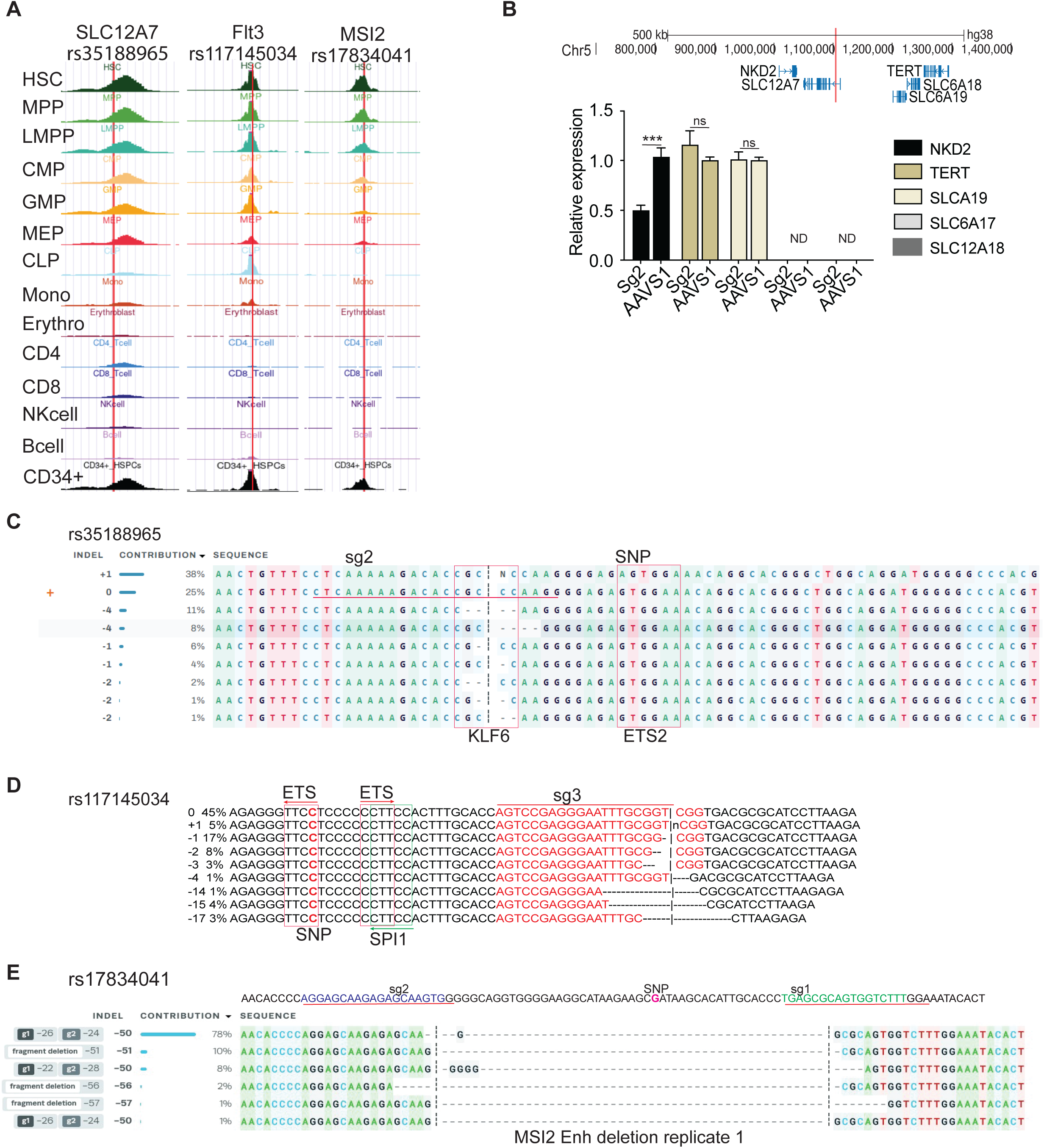
(A) ATAC peaks overlap of SLC12A7, Flt3 and MSI2 variants across 14 human hematopoietic mature and progenitor cell types. (B) Relative expression of genes located nearby SLC12A7 variant measured by qPCR. (C) Representative editing efficiency analysis of SLC12A7 variant containing enhancer in K562 cells using CRISPR technology as measured by ICE analysis on Sanger sequencing data. Positions of SNP and sgRNAs used are indicated (D) Representative editing efficiency analysis of FLT3 variant containing promoter in MOLM13 cells using CRISPR technology as measured by ICE analysis on Sanger sequencing data. Position of the SNP and sgRNAs are indicated in the sequencing data. (E) Representative editing efficiency of MSI2 variant containing enhancer microdeletion in K562 cells using CRISPR technology as measured by ICE analysis on Sanger sequencing data.

**Supplemental figure 3:**
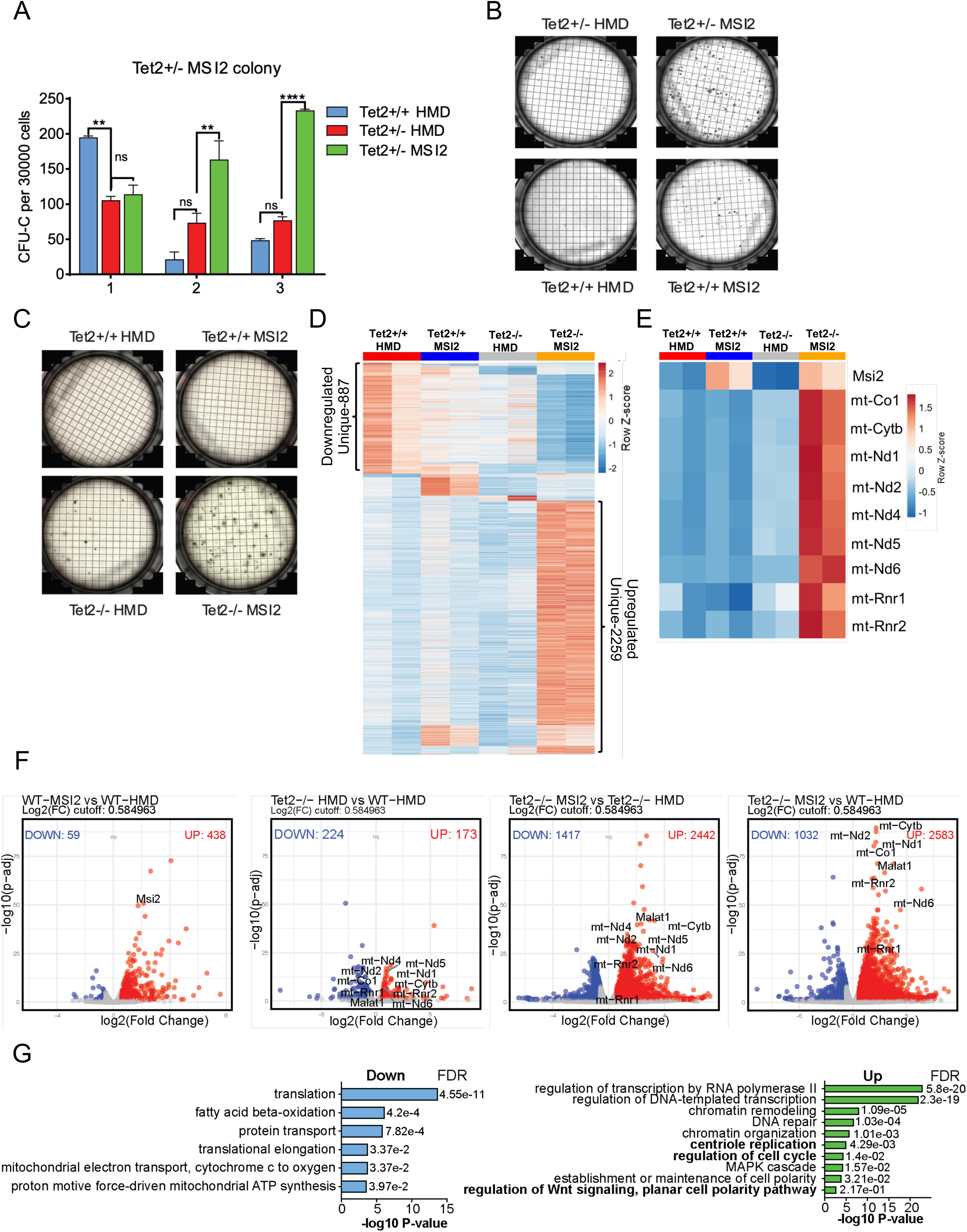
(A) CFU-C replating assay shows the colonies number from sorted murine GFP+ TET2+/- HSPCs. (B-C) Colony images of 2^nd^ plating from TET2+/- and TET2-/-HSPCs, respectively. (D) Heatmap showing expression of genes across all genotypes highlighting genes uniquely up- and down-regulated in the KO HMD group (E) Heatmap represents mitochondrial electron transport chain genes expression in different experimental groups. (F) Volcano plots show DEGs expression compared WT MSI2 vs WT HMD, Tet2-/- HMD vs WT HMD, Tet2-/- MSI2 vs Tet2-/- HMD, and Tet2-/- MSI2 vs WT HMD. (G) Gene ontology analysis of downregulated and upregulated pathways between Tet2^-/-^ MSI2 vs. WT HMD.

